# Autistic and non-autistic adults similarly experience statistical regularities

**DOI:** 10.64898/2026.07.09.737492

**Authors:** Kirsten Rittershofer, Emma K. Ward, Clare Press

## Abstract

Bayesian accounts of autism propose that perception is less influenced by prior expectations and more strongly driven by incoming sensory information in autistic than non-autistic individuals, with this altered balance cascading through the cognitive hierarchy to also influence higher cognitive functions. However, empirical support for these accounts remains mixed. Previous work has mostly tested these ideas in the context of objective environmental statistics, but recent work suggests that it may be subjective experience of structure, rather than structure itself, that shapes perceptual processing. Characterising these subjective experiences in autistic individuals is therefore crucial for understanding predictive processing in autism. In the present study, we thus examined subjective experience of statistical structure in autistic and non-autistic adults and tested how this experience relates to perceptual decisions. Participants were exposed to statistical regularities between action cues and visual stimuli (shapes), and we measured their speed and accuracy in reporting which shape they had seen. At the end of the study, participants were asked to estimate the probability and rate their surprise for each action-shape combination. Autistic and non-autistic participants showed similar subjective probability and surprise ratings and a comparable relationship between these ratings and perceptual decisions. Across participants, subjective ratings explained perceptual decisions better than objective structure. Together, these findings show that autistic and non-autistic adults experience statistical structure similarly, with these experiences exerting a similar influence on perceptual decisions – therefore suggesting that subjective experience plays a comparable role in predictive processing in autistic and non-autistic adults.

**Lay summary:** Our brains constantly learn patterns from past experience – for example, that dark clouds are often followed by rain – and use this knowledge to predict and interpret what we see. We studied how autistic and non-autistic adults experience these learned patterns by asking them how likely and surprising they found different events. We found that both groups gave similar ratings, and that these experiences shaped perception in similar ways.

## Introduction

Bayesian accounts of autism propose that perception is less influenced by prior expectations and more strongly driven by incoming sensory information in autistic than non-autistic individuals (Brock, 2012; Friston, Lawson, & Frith, 2013; Lawson, Rees, & Friston, 2014; Palmer, Lawson, & Hohwy, 2017; Pellicano & Burr, 2012; Sinha et al., 2014; van Boxtel & Lu, 2013; Van de Cruys, de-Wit, Evers, Boets, & Wagemans, 2013; Van de Cruys et al., 2014). Although these accounts differ in their precise proposed mechanisms – ranging from globally attenuated priors (Pellicano & Burr, 2012) to increased precision weighting of prediction errors (Lawson et al., 2014; Van de Cruys et al., 2014) – they converge on the suggestion that, at least sometimes, autistic perception relies more on sensory input and less on prior knowledge. This altered balance is proposed to affect all sensory processing and to cascade throughout the cognitive hierarchy, contributing to autistic characteristics across perceptual, social, and other domains.

Despite the appeal of these accounts as a unifying explanation of autism, empirical support remains mixed (Angeletos Chrysaitis & Seriès, 2023; Cannon, O’Brien, Bungert, & Sinha, 2021; Nobel Norrman et al., 2026; Sapey-Triomphe, Pattyn, Weilnhammer, Sterzer, & Wagemans, 2023). Some studies report findings consistent with an altered balance between prior expectations and sensory input, such as larger prediction error responses in autistic participants (van Laarhoven, Stekelenburg, Eussen, & Vroomen, 2020; also see Bravo et al., 2017; Ewbank et al., 2015; Lin et al., 2015; Manning, Tibber, Charman, Dakin, & Pellicano, 2015; Palmer, Paton, Kirkovski, Enticott, & Hohwy, 2015; Pomè, Wiesing, & Zimmermann, 2026; Pomè & Zimmermann, 2024, 2025; Pultsina et al., 2025; Sapey-Triomphe, Weilnhammer, & Wagemans, 2022; Skewes, Jegindø, & Gebauer, 2015; Turi et al., 2015; van Laarhoven, Stekelenburg, Eussen, & Vroomen, 2019; Zaidel, Goin-Kochel, & Angelaki, 2015). However, a growing body of work finds little to no evidence of differences in the relative weightings of priors and inputs (Achermann, Falck-Ytter, Bölte, & Nyström, 2021; Angeletos Chrysaitis, Jardri, Denève, & Seriès, 2021; Bosch, Fritsche, Utzerath, Buitelaar, & de Lange, 2022; Braukmann et al., 2018; Croydon, Karaminis, Neil, Burr, & Pellicano, 2017; Fazioli, Hadad, Denison, & Yashar, 2025b; Goris et al., 2021; Manning, Kilner, Neil, Karaminis, & Pellicano, 2017; Serences, Ester, Vogel, & Awh, 2009; Van de Cruys, Vanmarcke, Van de Put, & Wagemans, 2018; Van der Hallen, Vanmarcke, Noens, & Wagemans, 2017; Ward, Braukmann, Buitelaar, & Hunnius, 2020; Ward et al., 2021; Ward, Buitelaar, & Hunnius, 2022, 2024; Westner, Bosch, Utzerath, Buitelaar, & de Lange, 2025), suggesting that the predictive differences proposed by Bayesian accounts may not be as pervasive or robust as initially assumed.

Previous work on predictive processing in autism has mostly examined differences in predictions and prediction errors derived from objective environmental statistics. However, this approach largely overlooks the fact that we often have reportable subjective experiences concerning the structure of our environment, which frequently diverge from the objective statistical structure (Foster & Keane, 2015; Maguire & Maguire, 2009; Tversky & Kahneman, 1983) and may also shape predictive processes. Indeed, recent work of ours has shown that subjective experiences of structure explain more variance in perceptual decisions than the objective structure itself (Clarke, Rittershofer, Ward, Yon, & Press, 2026; also see Khoudary, Bornstein, & Peters, 2025). If perception is shaped by subjective experience of structure, rather than structure itself, characterising the nature of these subjective representations is crucial for an understanding of predictive processing in autism. In particular, determining whether autistic and non-autistic individuals differ in their subjective representations of structure, and whether these representations relate differently to perception, may clarify whether subjective experience contributes to proposed differences in predictive processing or whether any differences are more likely to arise from other mechanisms.

To address these questions, we recruited two groups of participants: one reporting a diagnosis of autism and one without. Participants were exposed to statistical regularities between action cues and visual stimuli (shapes), such that the shapes followed each action with different objective probabilities (ranging from 71% to 2%). On each trial, participants identified the presented shape, allowing us to assess the speed and accuracy of their perceptual decisions. At the end of the study, participants estimated the probability of each action-shape combination and rated how surprising they found each combination. We then compared subjective probability and surprise ratings between autistic and non-autistic individuals, and examined whether they were similarly related to perceptual decisions (reaction times and accuracy of the shape judgements) in the two groups. In addition to these group comparisons, we also tested whether any of these relationships varied as a function of individual differences in autistic traits, anxiety, or intolerance of uncertainty (IU).

## Methods

### Participants

Seventy participants per group (non-autistic group: 32 females, 38 males; mean age = 31.14 years, *SD* = 9.38; autistic group: 30 females, 40 males; mean age = 31.40 years, *SD* = 9.27) completed the study and were included in the analysis. Participants were recruited via Prolific (www.prolific.com) and reported normal or corrected-to-normal vision. Participants for the two groups were selected based on how they answered Prolific’s autism screening question when signing up *(‘Have you received a formal clinical diagnosis of autism spectrum disorder, made by a psychiatrist, psychologist, or other qualified medical specialist? This includes Asperger’s syndrome, Autism Disorder, High Functioning Autism or Pervasive Developmental Disorder.’*). Participants in the non-autistic group answered *‘No’* and participants in the autistic group responded with either *‘Yes – as a child’* or *‘Yes – as an adult’*. This question was repeated at the end of our study. Where participants’ responses differed from those provided to the Prolific screening question, we contacted them via Prolific to clarify their diagnosis status. Participants whose follow-up responses indicated that they did not meet the criteria for their assigned group, as well as those who did not respond to the follow-up request, were excluded from the analysis (nine from the non-autistic group and 13 from the autistic group). One additional participant was excluded due to a data-saving error. All participants gave informed consent prior to participation and were paid £5.35, equivalent to an hourly wage of approximately £8. The study complied with all relevant ethical regulations and was approved by the Research Department of Experimental Psychology Ethics Chair at UCL. The study was pre-registered prior to data collection at https://doi.org/10.17605/OSF.IO/QKJZ9.

### Procedure

The experiment was run online using Gorilla (www.gorilla.sc) and consisted of two parts. In the first part, participants completed online versions of three questionnaires: the GAD-7 Generalised Anxiety Scale (Spitzer, Kroenke, Williams, & Löwe, 2006), the IUS-12 Intolerance of Uncertainty Scale (Carleton, Norton, & Asmundson, 2007), and the RAADS-14 Autism Screener (Eriksson, Andersen, & Bejerot, 2013).

After 24 hours, participants were invited to complete the second part of the study. In this part, they were exposed to statistical regularities between action cues and visual stimuli (shapes). The objective probability of a particular shape following a given action varied across four levels (71%, 48%, 25%, and 2%; Figure 1A) and participants had to identify the presented shape. Each trial began with a fixation cross (500 ms), followed by the word *‘index’*, *‘middle’*, or *‘ring’*, specifying which action to perform – a key press with the corresponding finger of their right hand. Once participants performed the correct action, a shape (circle, triangle, square, or hexagon) appeared on the screen, and their task was to indicate as quickly and accurately as possible which shape was shown. The shape remained on the screen for 400 ms, or less if they responded before that. Participants responded using their left hand, pressing *‘1’* to *‘4’* for circle, triangle, square, and hexagon, respectively, and there was no time limit for responses. After responding, a blank inter-trial interval of either 500 or 1000 ms (pseudo-randomised) followed before the next trial began (Figure 1B). Trial order within each block was randomised and participants received feedback on their accuracy and mean reaction time after each block. The experiment consisted of four blocks of 75 trials. Each participant was pseudo-randomly assigned to one of 20 different randomly generated action-shape mappings.

**Figure 1:**
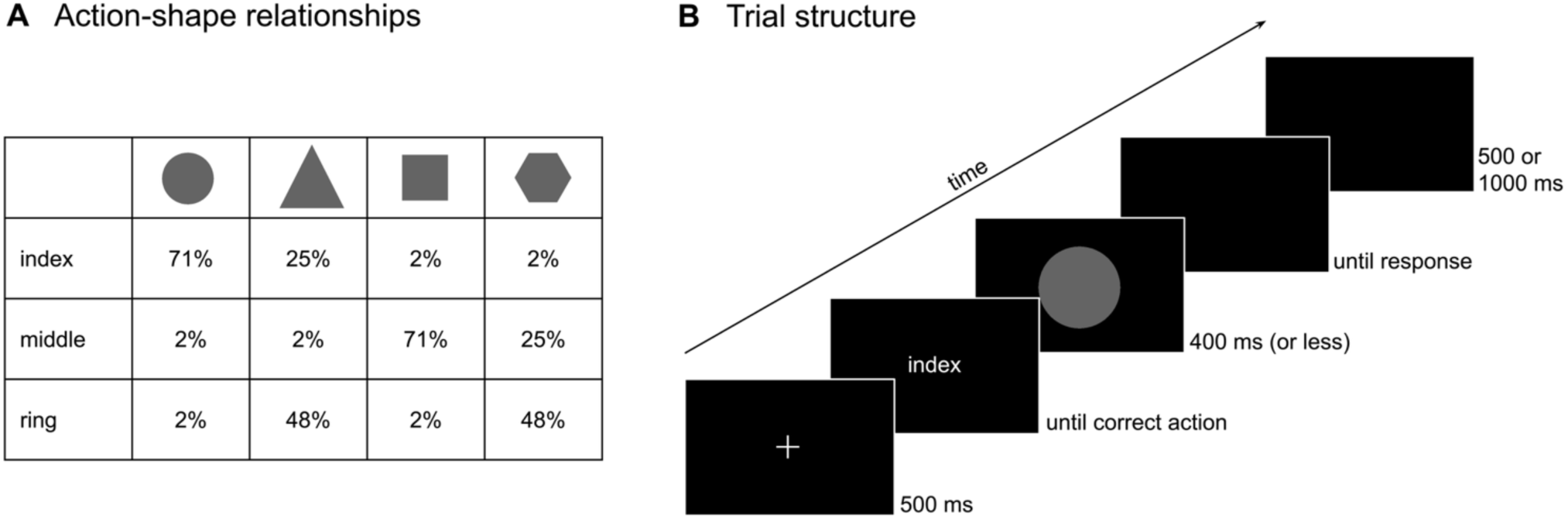
Design. **(A)** Relationships between actions and shapes with the objective probability of a particular shape following a given action ranging from 71% to 48%, 25%, and 2%. At the end of the experiment, participants rated the surprise and probability for each of these 12 action-shape combinations. **(B)** Trial structure. On each trial, participants were cued to perform an action and then presented with an outcome – one of the four shapes according to the relationships outlined in (A). Participants’ task was to indicate as quickly and accurately as possible which shape was shown.

Following the task, participants were asked to rate how surprised they were by each action-shape combination on a scale from *‘0 (not surprised at all)’* to *‘100 (highly surprised)’* (e.g., *‘How surprised were you when after pressing down a key (‘H’) with your right index finger a circle appeared?*). They then also rated the probability of each outcome on a scale from *‘never (0%)’* to *‘always (100%)’* (e.g., *‘When you pressed down a key (‘H’) with your right index finger, how often was it followed by a circle?’*). For all ratings, the initial slider position was at the centre (50), and participants were required to move it away from and back to 50 if they wanted to select this value.

### Analysis

The data were analysed using linear mixed-effects models with the lme4 package in R (Bates, Mächler, Bolker, & Walker, 2015). Continuous predictors were z-scored prior to analysis, and categorical predictors were sum-to-zero contrast coded. Trials with reaction times (RTs) below 100 or above 2500 ms were excluded from all analyses. For RT analyses, only trials with correct responses were included.

For all analyses, we compared five models. A null model, in which the effect of interest was assumed to be constant across participants, was compared to models in which the predictors of interest interacted with either a categorical group variable (autistic, non-autistic) or with continuous scores on the autism (RAADS-14), anxiety (GAD-7), or intolerance of uncertainty (IUS-12) questionnaires. Model comparison was performed using the Bayesian Information Criterion (BIC), with lower values indicating better model fit. To aid interpretation, we also report approximate Bayes factors comparing each alternative model to the null model, derived from the BIC values following Masson (2011): BF_01_ ≈ e^ΔBIC/2^. Model comparison results using Akaike Information Criterion (AIC), which impose a less stringent penalty on complexity, are reported in the Supplementary Materials, but yielded similar conclusions to the BIC comparisons.

To test whether subjective probability and surprise ratings tracked objective probability, and whether these relationships varied according to group membership or questionnaire scores, we first specified the following null model:

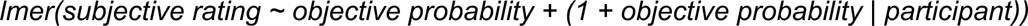

This model was then compared to models in which the fixed effect of objective probability interacted with either a categorical group variable or one of the continuous questionnaire scores.

To examine whether perceptual decisions about the shapes were influenced by objective probability and subjective ratings, and whether these relationships differed according to group membership or questionnaire scores, we specified the following null models. In these models, reaction times (RTs) and accuracy were predicted from objective probability, rating (either subjective surprise or subjective probability), and trial number, which was included to account for global changes in performance over the course of the experiment due to practice or fatigue. For RT analyses, linear mixed-effects models predicted log-transformed RTs, whereas for accuracy analyses, logistic mixed-effects models predicted correct versus incorrect responses. The null models were specified as follows:

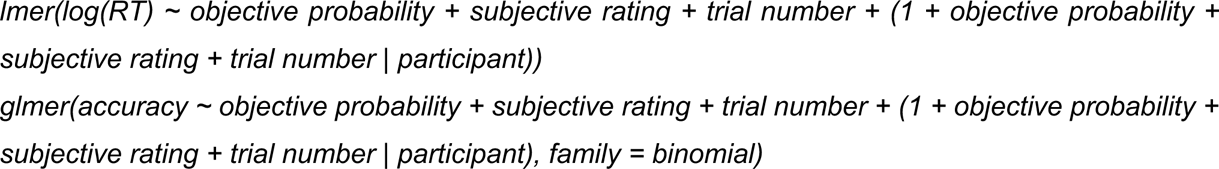

These null models were then compared to models in which the fixed effects of objective probability and subjective rating interacted with either a categorical group variable or one of the continuous questionnaire scores.

## Results

### Questionnaires

Participants in the autistic group scored significantly higher on all three questionnaires than those in the non-autistic group. On the RAADS-14 autism screener (possible score range: 0-42), the autistic group (*M* = 22.74, *SD* = 10.57) scored significantly higher than the non-autistic group (*M* = 11.37, *SD* = 8.04; *t*(138) = −7.17, *p* < .001, *d* = −1.21; Figure 2A). The RAADS-14 is composed of three subscales, and the autistic group also scored significantly higher on each subscale. For the mentalizing deficits subscale (possible score range: 0-21), the autistic group had a higher mean score of 11.83 (*SD* = 6.25), compared to the non-autistic group’s mean of 5.54 (*SD* = 4.62; *t*(138) = −6.77, *p* < .001, *d* = −1.14; Figure 2D). On the social anxiety subscale (possible score range: 0-12), the autistic group (*M* = 6.19, *SD* = 3.42) also scored significantly higher than the non-autistic group (*M* = 3.60, *SD* = 3.47; *t*(138) = −4.44, *p* < .001, *d* = −0.75; Figure 2E). Similarly, on the sensory reactivity subscale (possible score range: 0-9), the autistic group (*M* = 4.73, *SD* = 2.87) scored significantly higher compared to the non-autistic group (*M* = 2.23, *SD* = 2.13; *t*(138) = −5.85, *p* < .001, *d* = −0.99; Figure 2F). These consistently higher scores on the overall RAADS-14 and its subscales in the autistic group confirm that our group classification captured meaningful differences, supporting the validity of our autistic versus non-autistic group comparisons. In addition, we also conducted all analyses with RAADS-14 score as a continuous predictor. Participants in the autistic group also scored significantly higher on the GAD-7 generalised anxiety scale (possible score range: 0-21). The autistic group had a mean score of 8.39 (*SD* = 5.62), compared to 5.96 (*SD* = 5.14) in the non-autistic group (*t*(138) = −2.67, *p* = .009, *d* = −0.45; Figure 2B). Similarly, on the IUS-12 intolerance of uncertainty scale (possible score range: 12-60), the autistic group scored significantly higher (*M* = 39.00, *SD* = 10.06) than the non-autistic group (*M* = 32.59, *SD* = 11.03; *t*(138) = −3.59, *p* < .001, *d* = −0.61; Figure 2C).

**Figure 2:**
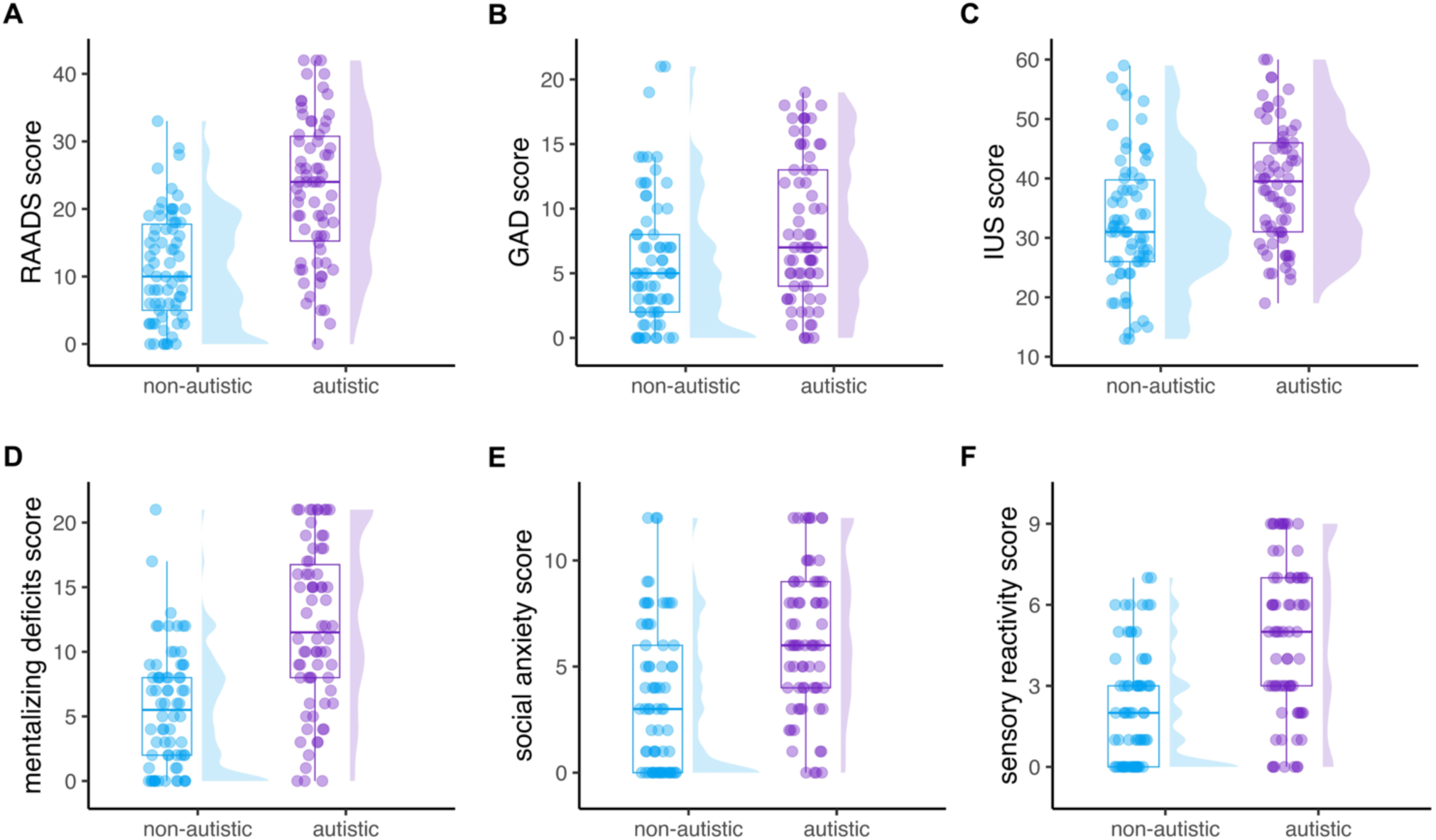
Questionnaire results. **(A)** RAADS-14 autism scores (0-42 range). The autistic group scored significantly higher than the non-autistic group. **(B)** GAD-7 generalised anxiety scores (0-21 range) were significantly higher in the autistic group. **(C)** IUS-12 intolerance of uncertainty scores (12- 60 range) also showed significantly higher values in the autistic group. **(D)** Mentalizing deficits subscale of the RAADS-14 (0-21 range), with significantly higher scores in the autistic group. **(E)** Social anxiety subscale scores (0-12 range) were significantly higher in the autistic group. **(F)** Sensory reactivity subscale scores (0-9 range) were also significantly higher in the autistic group. Boxes denote lower, middle, and upper quartiles, and whiskers extend to 1.5 times the interquartile range. Coloured dots represent individual participant data. The half-violin plots display probability density estimates of the scores.

### Ratings

To examine whether ratings of subjective surprise and subjective probability varied as a function of group or participants’ scores on the autism (RAADS-14), anxiety (GAD-7), or intolerance of uncertainty questionnaires (IUS-12), we compared five models. In all models, objective outcome probability was included as a predictor of the ratings. The null model assumed that this effect was constant across participants, whereas the alternative models included interactions between objective probability and either a categorical group variable or continuous scores on one of the three questionnaires. Model comparison was then used to determine which model provided the best fit to the data. To aid interpretation, we additionally report Bayes factors (BF_01_), quantifying the evidence in favour of the null model relative to each alternative model.

For the subjective surprise ratings, the null model best fit the data (BIC = 15339.68), outperforming the models including interactions with group (BIC = 15354.47, BF_01_ = 1633.97), autism scores (BIC = 15354.39, BF_01_ = 1568.38), anxiety scores (BIC = 15354.46, BF_01_ = 1621.94), or intolerance of uncertainty scores (BIC = 15354.09, BF_01_ = 1347.57). The null model revealed a significant negative effect of objective probability on the subjective surprise ratings (*β* = -5.449, 95% CI [-7.151, -3.747], *p* < .001), indicating that objectively less probable outcomes were rated as more surprising by the participants. Importantly, however, this relationship did not vary by group, nor was it modulated by individual differences in autistic traits, anxiety, or intolerance of uncertainty (Figure 3A).

**Figure 3:**
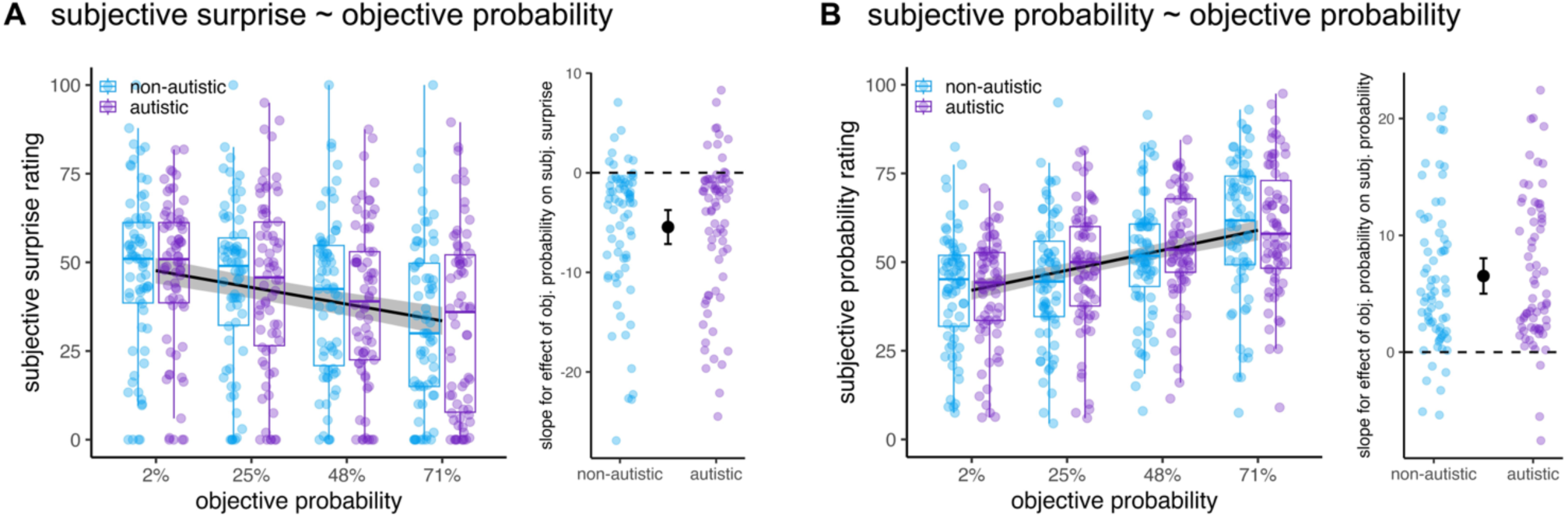
Rating results. **(A)** Subjective surprise ratings and **(B)** subjective probability ratings. Left panels show participants’ mean ratings across the four levels of objective probability for the autistic (purple) and non-autistic (blue) groups, alongside the fixed-effect fit from the winning null models (i.e., excluding group membership). Surprise ratings decreased and subjective probability ratings increased with objective probability. Neither relationship was affected by group membership or questionnaire scores. Right panels show participant-specific slopes for the effect of objective probability on each rating, derived from the winning null models and displayed separately by group for visualisation. Slopes were predominantly negative for surprise ratings and positive for subjective probability ratings. In the left panels, coloured dots represent each participant’s mean rating. Boxes denote lower, middle, and upper quartiles, and whiskers extend to 1.5 times the interquartile range. Black lines and shaded ribbons represent the fixed-effect estimates from the winning null models and their 95% confidence intervals (CIs). In the right panels, coloured dots represent participant-specific slopes, and black points and error bars indicate the fixed-effect slope estimates and their 95% CIs. Dashed horizontal lines at zero indicate no effect of objective probability on the ratings.

A similar pattern was observed for the subjective probability ratings. The null model again fit the data best (BIC = 14906.14), outperforming the models including interactions with group (BIC = 14920.42, BF_01_ = 1262.35), autism (BIC = 14920.05, BF_01_ = 1048.73), anxiety (BIC = 14918.60, BF_01_ = 507.82), or intolerance of uncertainty scores (BIC = 14918.55, BF_01_ = 495.64). The null model revealed a significant positive effect of objective probability on the subjective probability ratings (*β* = 6.518, 95% CI [5.003, 8.034], *p* < .001), such that objectively more probable outcomes were also rated as more probable by the participants. As with the subjective surprise ratings, this relationship did not vary by group and was not modulated by questionnaire scores either (Figure 3B).

### Reaction times

We next examined whether reaction times in identifying the presented shape were influenced by objective outcome probability and participants’ subjective ratings, and whether these effects differed as a function of group or questionnaire scores. To do so, we compared models predicting reactions times from objective probability and a rating (either subjective surprise or subjective probability). A null model assuming constant effects across participants was compared to models including interactions with group or questionnaire scores.

For the models including objective probability and subjective surprise rating, the null model (BIC = 8722.78) outperformed models in which the two predictors interacted with group (BIC = 8751.69, BF_01_ > 10^6^), autism (BIC = 8752.51, BF_01_ > 10^6^), anxiety (BIC = 8751.66, BF_01_ > 10^6^), or intolerance of uncertainty scores (BIC = 8750.49, BF_01_ = 10^6^). In this model, objective probability had a significant negative effect on reaction times (*β* = -0.012, 95% CI [-0.021, -0.003], *p* = .009), indicating faster responses for objectively more probable outcomes. Subjective surprise showed a significant positive effect (*β* = 0.043, 95% CI [0.023, 0.064], *p* < .001), with slower responses to outcomes rated as more surprising. Importantly, these effects did not differ by group and were not modulated by questionnaire scores (Figure 4A).

**Figure 4:**
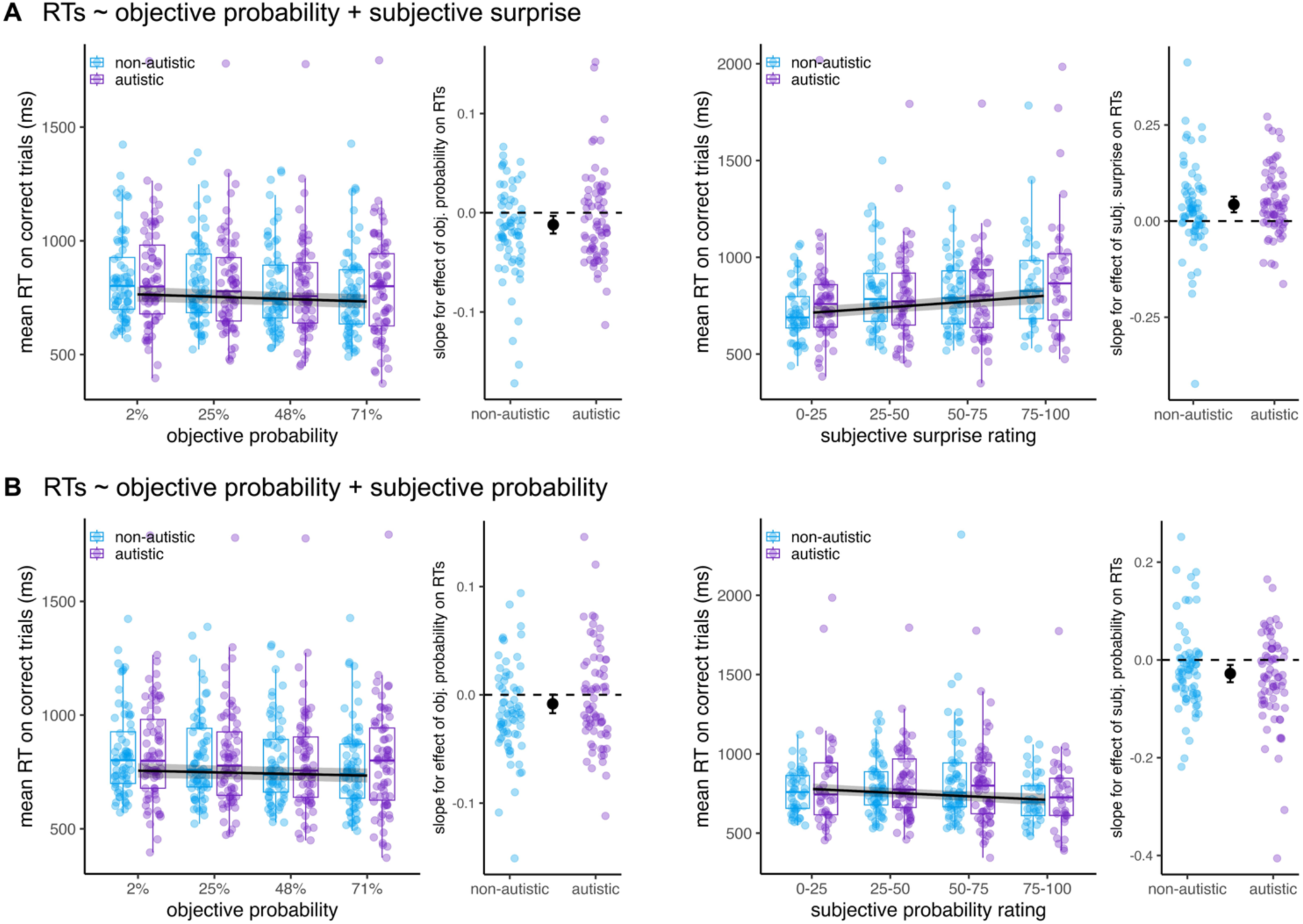
Reaction time results. **(A)** RTs predicted from objective probability and subjective surprise. RTs decreased with increasing objective probability, with predominantly negative participant-specific slopes (left two plots), and increased with subjective surprise, with predominantly positive slopes (right two plots). **(B)** RTs predicted from objective and subjective probability. When included alongside subjective probability, objective probability no longer significantly predicted RTs, with participant-specific slopes centred around zero (left two plots), whereas RTs decreased with increasing subjective probability, with predominantly negative slopes (right two plots). None of these relationships was affected by group membership or questionnaire scores. The RT plots show each participant’s mean RT on correct trials (coloured dots) either across the four levels of objective probability or within four equal-width bins of subjective ratings (created for visualisation purposes only). The slope plots show the corresponding participant-specific slopes derived from the winning null models. Black lines (RT plots) and black points (slope plots) show the fixed-effect estimates from the winning null models. Other plotting conventions are as in Figure 3.

For the models including objective probability and subjective probability rating, the null model (BIC = 9030.95) outperformed models in which the two predictors interacted with group (BIC = 9058.88, BF_01_ > 10^6^), autism (BIC = 9062.50, BF_01_ > 10^6^), anxiety (BIC = 9061.07, BF_01_ > 10^6^), or intolerance of uncertainty scores (BIC = 9059.23, BF_01_ > 10^6^). In this model, subjective probability had a significant negative effect on reaction times (*β* = -0.028, 95% CI [-0.046, -0.010], *p* = .003), indicating faster responses for outcomes rated as more probable. In contrast, objective probability did not significantly predict reaction times (*β* = -0.008, 95% CI [-0.017, 0.0003], *p* = .061). Again, these effects did not differ by group and were not modulated by questionnaire scores (Figure 4B).

### Accuracy

As we did for reaction times, we examined whether participants’ accuracy in identifying the presented shape was influenced by objective outcome probability and subjective ratings, and whether these effects differed as a function of group or questionnaire scores. To do so, we compared models predicting accuracy from objective probability and rating (either subjective surprise or subjective probability). A null model assuming constant effects across participants was again compared to models including interactions with group or questionnaire scores.

For the models including objective probability and subjective surprise rating, the null model (BIC = 18823.35) outperformed models in which the two predictors interacted with group (BIC = 18854.50, BF_01_ > 10^6^), autism (BIC = 18854.95, BF_01_ > 10^6^), anxiety (BIC = 18851.72, BF_01_ > 10^6^), or intolerance of uncertainty scores (BIC = 18855.02, BF_01_ > 10^6^). In this model, subjective surprise had a significant negative effect on accuracy (*β* = -0.127, 95% CI [-0.240, -0.013], *p* = .029), indicating lower accuracy for outcomes rated as more surprising. In contrast, objective probability did not significantly predict accuracy (*β* = 0.021, 95% CI [-0.050, 0.092], *p* = .561). These effects did not differ by group and were not modulated by questionnaire scores (Figure 5A).

**Figure 5:**
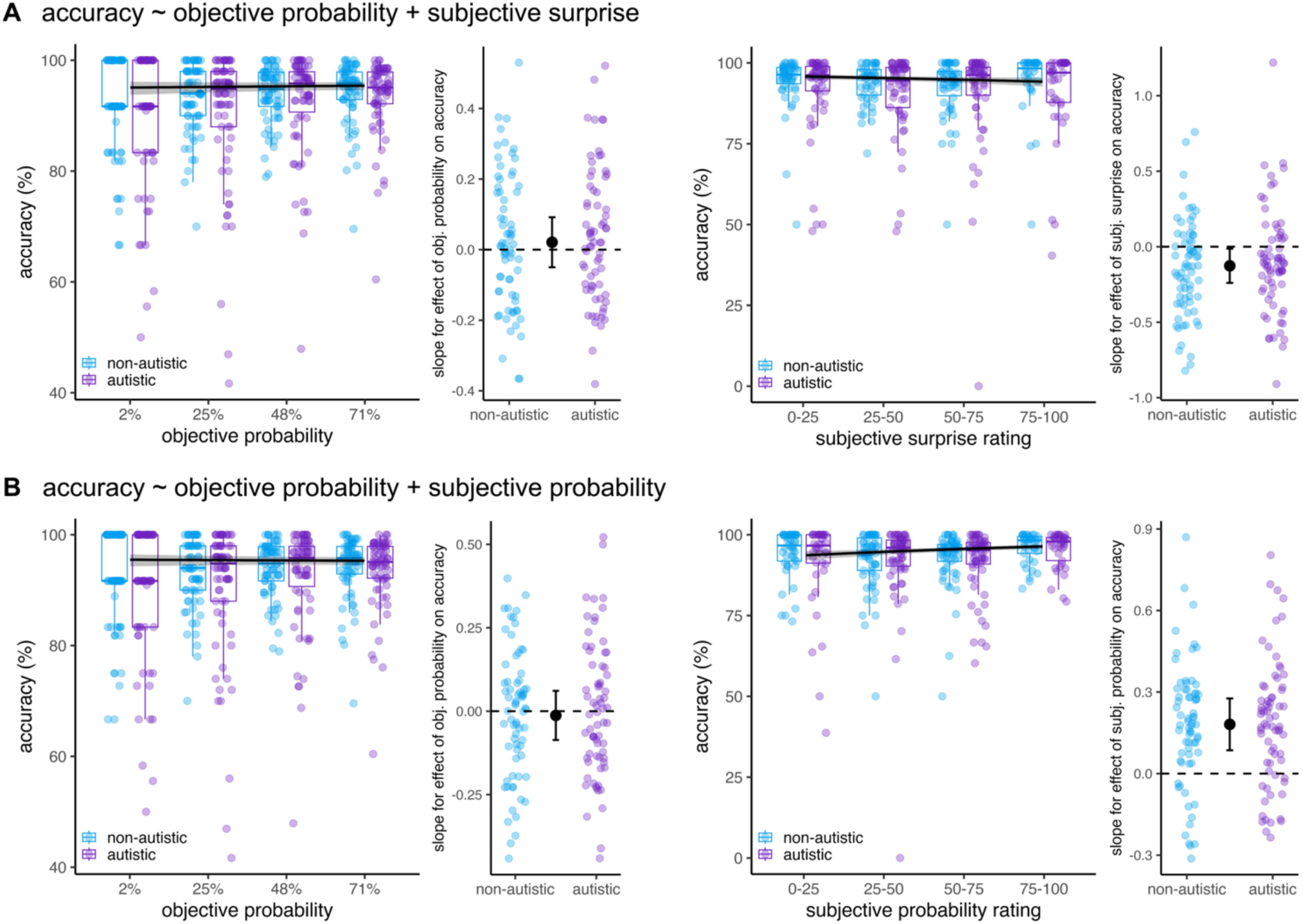
Accuracy results. **(A)** Accuracy predicted from objective probability and subjective surprise. Accuracy did not change as a function of objective probability, with participant-specific slopes centred around zero (left two plots), but decreased with higher subjective surprise, with predominantly negative slopes (right two plots). **(B)** Accuracy predicted from objective and subjective probability. Objective probability had no effect on accuracy, with participant-specific slopes centred around (left two plots), but accuracy increased with higher subjective probability, with predominantly positive slopes (right two plots). None of these relationships was affected by group membership or questionnaire scores. The accuracy plots show each participant’s accuracy (coloured dots) either across the four levels of objective probability or within four equal-width bins of subjective ratings (created for visualisation purposes only). The slope plots show the corresponding participant-specific slopes derived from the winning null models. Black lines (RT plots) and black points (slope plots) show the fixed-effect estimates from the winning null models. Other plotting conventions are as in Figure 3.

For the models including objective probability and subjective probability rating, the null model (BIC = 18831.72) again provided the best fit, outperforming models including interactions with group (BIC = 18863.00, BF_01_ > 10^6^), autism (BIC = 18862.98, BF_01_ > 10^6^), anxiety (BIC = 18860.73, BF_01_ > 10^6^), or intolerance of uncertainty scores (BIC = 18863.25, BF_01_ > 10^6^). In this model, subjective probability had a significant positive effect on accuracy (*β* = 0.181, 95% CI [0.086, 0.276], *p* < .001), indicating higher accuracy for outcomes rated as more probable. Objective probability did not significantly predict accuracy (*β* = -0.013, 95% CI [-0.086, 0.061], *p* = .735). As above, these effects did not differ by group and were not modulated by questionnaire scores (Figure 5B).

## Discussion

In the present study, we investigated how autistic adults experience statistical structure, and how this experience shapes perceptual decisions. Following a probabilistic cue-outcome task, participants estimated the probability of each cue-outcome combination and rated how surprising they found each combination. Autistic and non-autistic participants gave similar subjective probability and surprise ratings, and these ratings showed comparable relationships with the speed and accuracy of their perceptual decisions, explaining behaviour better than objective structure across participants. Likewise, neither subjective ratings nor their relationship with perceptual decisions varied as a function of autistic traits, anxiety, or intolerance of uncertainty.

These findings suggest that autistic and non-autistic individuals experience statistical structure similarly, and that these experiences influence perceptual decisions in comparable ways. Given recent work showing that perceptual decisions are more strongly influenced by subjective experience of structure than objective structure itself (Clarke et al., 2026; also see Khoudary et al., 2025), characterising these subjective experiences is crucial for understanding predictive processing in autism and can shed light on the mixed findings in the literature. Specifically, if autistic individuals differed in their subjective experience of structure, or in how this experience relates to perception, group differences would be expected to primarily emerge in paradigms where subjective experience plays a stronger role in shaping perception. However, the similar patterns observed in both groups render this explanation unlikely, suggesting instead that the mixed findings are more likely to reflect variation in other mechanisms.

While we observed comparable subjective representations of statistical structure at the end of learning, this does not necessarily imply that autistic and non-autistic individuals acquired these representations in the same way. Indeed, a recent review concluded that several studies provide evidence for differences in learning between autistic and non-autistic individuals (Angeletos Chrysaitis & Seriès, 2023; but also see Pultsina et al., 2025). Such differences may not have been captured in the present study because subjective probability and surprise were measured only once, at the end of learning. It is therefore possible that group differences emerge during the acquisition of statistical regularities but become less apparent once stable representations have formed. Consistent with this possibility, Solomon, Smith, Frank, Ly, and Carter (2011) reported group differences during early learning that were no longer evident by the end of training. Future studies could test this possibility by measuring subjective experience throughout learning, rather than only at the end of learning. Building on trial-by-trial prediction measures in autism (Sapey-Triomphe et al., 2022), future studies could obtain repeated ratings of subjective probability and surprise, as in some of our previous work (Clarke et al., 2026), to track how subjective experience evolves as autistic and non-autistic individuals acquire statistical regularities.

An alternative possibility is that autistic and non-autistic individuals do not differ in their ability to learn statistical regularities per se, but rather in meta-learning processes – that is, in judging when to learn. This involves estimating whether changes in the environment reflect genuine shifts in the underlying statistical structure or merely random fluctuations. This suggestion is consistent with emerging evidence for differences in other second-order cognitive processes, specifically metacognition (i.e., judging when we know; Fazioli, Hadad, Denison, & Yashar, 2025a; Johnstone, Friston, Rees, & Lawson, 2022). It is also consistent with findings from volatile environments, where statistical relationships change over time and learners must continually assess whether previously learned relationships remain reliable. Studies have reported difficulties learning under volatile conditions (Robic et al., 2015; Sevgi, Diaconescu, Henco, Tittgemeyer, & Schilbach, 2020), altered brain activation in regions associated with uncertainty estimation (Nobel Norrman et al., 2026), reduced flexibility in adjusting predictions in a changing environment (Sapey-Triomphe et al., 2022), and a tendency to overestimate environmental volatility (Lawson, Mathys, & Rees, 2017), although the evidence remains mixed (Goris et al., 2021; Manning et al., 2017). By contrast, although participants in the present study had to learn novel cue-outcome relationships, these relationships remained stable throughout the experiment and therefore placed relatively little demand on deciding when updating was required. Future work using a similar paradigm in which cue-outcome relationships change over time could test whether subjective experience differs under conditions that place greater demands on meta-learning.

Participants were assigned to groups based on self-reported autism diagnoses provided when signing up on Prolific, and we were unable to independently verify these reports. Although we excluded participants who gave inconsistent responses across repeated questions, some degree of misclassification cannot be ruled out. Nevertheless, the groups differed significantly on the RAADS-14 autism screener, including all three subscales (mentalizing deficits, social anxiety, and sensory reactivity), indicating that the group assignment captured meaningful variation in autistic characteristics. Moreover, analyses using continuous questionnaire scores, rather than categorical group membership, likewise revealed no evidence that subjective experience or its relationship with perceptual decisions varied as a function of autistic traits. Together, these findings suggest that the absence of group differences is unlikely to be attributable to the method of participant recruitment and group assignment.

To conclude, in the present study we examined autistic adults’ subjective experience of environmental structure and how this experience, as well as objective structure itself, influences perceptual decisions. We found similar subjective experiences of statistical structure, as well as comparable influences on perceptual decisions, in autistic and non-autistic adults, with subjective experience of structure influencing decisions to a greater extent than the objective structure itself. These findings were also consistent across individual differences in autistic traits, anxiety, and intolerance of uncertainty. Together, these findings suggest that subjective experience contributes similarly to predictive processing in autistic and non-autistic adults, making it unlikely that variation in subjective experience accounts for the mixed predictive processing findings across the literature and pointing instead towards variation in other mechanisms.

## Acknowledgements

This work was funded by a European Research Council (ERC) consolidator grant (101001592) under the European Union’s Horizon 2020 research and innovation programme, and a Leverhulme Trust project grant (RPG-2022-358), both awarded to CP. We are grateful to other members of the Action and Perception Lab for helpful discussions throughout the project.

## Data availability statement

All data and analysis code required to reproduce the findings reported in this study will be made publicly available on the Open Science Framework (OSF) upon publication. The experimental materials, including the stimuli and Gorilla task spreadsheets, will also be made available.

