## Supplementary Materials for "Autistic and non-autistic adults similarly experience statistical regularities"

### Model comparisons using AIC

**Ratings.** For the subjective surprise ratings, the null model best fit the data (AIC = 15307.12), outperforming the models including interactions with group (AIC = 15311.07), autism scores (AIC = 15310.99), anxiety scores (AIC = 15311.06), or intolerance of uncertainty scores (AIC = 15310.69). For the subjective probability ratings, the null model again fit the data best (AIC = 14873.58), outperforming the models including interactions with group (AIC = 14877.02), autism (AIC = 14876.64), anxiety (AIC = 14875.19), or intolerance of uncertainty scores (AIC = 14875.15).

**Reaction times.** For the models including objective probability and subjective surprise rating, the null model (AIC = 8594.31) outperformed models in which the two predictors interacted with group (AIC = 8597.52), autism (AIC = 8598.34), anxiety (AIC = 8597.50), or intolerance of uncertainty scores (AIC = 8596.32). For the models including objective probability and subjective probability rating, the null model (AIC = 8902.48) outperformed models in which the two predictors interacted with group (AIC = 8904.72), autism (AIC = 8908.33), anxiety (AIC = 8906.91), or intolerance of uncertainty scores (AIC = 8905.06).

**Accuracy.** For the models including objective probability and subjective surprise rating, the null model (AIC = 18702.46) outperformed models in which the two predictors interacted with group (AIC = 18707.71), autism (AIC = 18708.16), anxiety (AIC = 18704.93), or intolerance of uncertainty scores (AIC = 18708.23). For the models including objective probability and subjective probability rating, the null model (AIC = 18710.83) again provided the best fit, outperforming models including interactions with group (AIC = 18716.21), autism (AIC = 18716.19), anxiety (AIC = 18713.94), or intolerance of uncertainty scores (AIC = 18716.46).

### Deviations from the pre-registration

There were some minor deviations from the pre-registered analysis plan. For the ratings, reaction times, and accuracy analyses, we compared five models rather than the three originally specified. As pre-registered, we compared a null and a model including interactions with group membership. However, instead of fitting a single model containing interactions with all three questionnaire scores, we fitted three separate models, each including interactions with a single questionnaire score. This modification was made to allow a fairer comparison between models. A model containing interactions with all three questionnaire scores is substantially more complex than the group model and is therefore penalized more heavily by information criteria such as BIC and AIC. Fitting separate questionnaire models ensured that each alternative model had the same level of complexity, allowing differences in model fit to be more directly attributed to the predictor of interest rather than differences in model complexity.

For the reaction times and accuracy analyses, we pre-registered models that included objective probability, subjective surprise, and subjective probability as simultaneous predictors. In the final analyses, we instead fitted separate models for subjective surprise and subjective probability, with each model including objective probability together with one of these subjective measures. This modification reflects the fact that subjective surprise and subjective probability are closely related measures of participants' subjective experience of statistical structure, and our primary aim was not to compare their unique contributions, but rather to examine how subjective experience (measured in two different ways) relates to perceptual decisions beyond objective probability. This approach is also consistent with our previous work on subjective experience (Clarke, Rittershofer, Ward, Yon, & Press, 2026).

Nevertheless, if conducting the pre-registered analyses, the primary conclusions reported in the main text remain unaltered.
